# A nearly complete and phased genome assembly of a Colombian *Trypanosoma cruzi* TcI strain and the evolution of gene families

**DOI:** 10.1101/2023.07.17.549441

**Authors:** Maria Camila Hoyos Sanchez, Hader Sebastian Ospina Zapata, Brayhan Dario Suarez, Carlos Ospina, Hamilton Julian Barbosa, Julio Cesar Carranza Martinez, Gustavo Adolfo Vallejo, Daniel Urrea Montes, Jorge Duitama

## Abstract

Chagas is an endemic disease in tropical regions of Latin America, caused by the parasite *Trypanosoma cruzi*. High intraspecies variability and genome complexity have been challenges for the development of genomic variation databases, needed to conduct studies in evolution, population genomics, and identification of genomic elements related to virulence and drug resistance in *T. cruzi*. Here we present a chromosome-level phased assembly of a *T. cruzi* strain (Dm25), isolated from a reservoir of the species *Didelphis marsupialis* located at the Tolima department in Colombia, and belonging to the TcI DTU. We obtained a primary haplotype composed of 32 chromosomes, 30 of them assembled in a single contig, and one complete copy of the maxicircle. While 29 chromosomes show a large collinearity with the assembly of the Brazil A4 strain, three chromosomes with a high density of repeat elements show a large divergence, compared to the Brazil A4 assembly. Considering that the distribution of heterozygous sites suggest that Dm25 is diploid, we assembled a second haplotype for 31 chromosomes, achieving an average of three contigs per chromosome. Nucleotide and protein evolution statistics indicate that *T. cruzi* Marinkellei separated before the diversification of *T. cruzi* in the known DTUs. Interchromosomal paralogs of dispersed gene families and histones appeared before but at the same time have a more strict purifying selection, compared to other repeat families. Previously unreported large tandem arrays of protein kinases and histones were identified in this assembly. Over one million variants obtained from Illumina reads aligned to the primary assembly clearly separate the main DTUs. We expect that this new assembly will be a valuable resource for further studies on evolution and functional genomics of *Trypanosomatids*.

## INTRODUCTION

Chagas disease (CD), also known as American trypanosomiasis, is a tropical disease caused by the protozoan parasite *Trypanosoma cruzi* that belongs to the Kinetoplastida class and the Trypanosomatidae family (Rassi, Rassi, & De Rezende, 2012). This disease is a public health problem, it is estimated that around 6 to 7 million people worldwide are infected with *T. cruzi* (WHO, 2020). About 30,000 new cases are registered each year, an average of 12,000 deaths, and 9,000 newborns are infected during pregnancy (OPS, n.d). Particularly in Colombia, a prevalence between 700,000-1,200,000 infected people and 8,000,000 individuals at risk of acquiring the infection has been estimated (MinSalud, 2010).

CD is found mainly in endemic areas of 21 Latin American countries, including Colombia. However, in recent decades it has spread to other countries such as the United States, Canada, and some European and African countries, due to the migration of the infected population (Schmunis & Yadon, 2010; WHO, 2020) or the presence of the vector and parasite (Curtis-Robles et al., 2018).

The life cycle of the parasite is complex since it has several forms of development in vectors and mammalian hosts, such as metacyclic trypomastigote (infective form), amastigote (intracellular form), epimastigotes (replicate form in the vector insect) (Echeverria & Morillo, 2019; Rassi et al., 2012). More than 150 species of domestic or wild mammals, such as dogs, cats, rodents, common opossum and armadillos, can be reservoirs of the parasite (Echeverria & Morillo, 2019; Rassi et al., 2012). In addition, about 152 species of triatomine insects are known and all have the potential to act as vectors of *T. cruzi* (De Oliveira et al., 2018).

*T. cruzi* is considered a parasite with a wide genetic diversity (Jiménez et al., 2019; Manoel-Caetano & Silvia, 2007; Zingales et al., 2018). This parasite has been classified into 7 different Discrete Typing Units (DTU) (TcI-TcVI and TcBat) (Tibayrenc, 1998; Sturm et al.,, 2003; Zingales et al., 2012; Barnabé, Mobarec, Jurado, Cortez, & Breniere, 2016; Marcili et al., 2009). Additionally, subdivisions or genotypes within some DTUs such as *T. cruzi* I (TcI) have been suggested (Cura et al., 2010; Falla et al., 2009; Gómez-Hernández et al., 2019; Herrera et al., 2007, Herrera et al., 2009; Llewellyn et al., 2009), which demonstrates the wide genetic variability of the parasite. The different DTUs present associations with transmission cycles, geography, vector species, and clinical manifestation to a certain extent (Hernández, Salazar, et al., 2016; Zingales et al., 2012). The variability of *T. cruzi* isolates circulating in Colombia and their association with the eco-epidemiology of Chagas disease have been studied for several years. The results show that the Colombian *T. cruzi* isolates present great genetic variability and suggest that TcI is predominant throughout the territory (Triana et al., 2006; Mejia-Jaramillo et al., 2009; Ramirez et al., 2013; Villa et al., 2013). Particularly, TcI has been associated with heart disease in chagasic patients in Colombia (Ramírez et al., 2010).

This extensive genetic variability is a result of the complexity of its genome. It has been widely reported that the genome of *T. cruzi* has extraordinary plasticity between strains, with a total length ranging between 40-140 Mb (Díaz-Viraqué et al., 2019; Souza et al., 2011). It is considered a generally diploid organism with the presence of aneuploidy in some hybrid strains (Minning, et al., 2011; Reis-Cunha et al., 2015; Souza et al., 2011). Its proteome has about 22,000 proteins (El-sayed et al., 2005). More than 50% of the genome consists of repetitive sequences, mainly represented by large multigene families encoding surface proteins, retrotransposons, telomeric repeats, and satellites (El-sayed et al., 2005; Reis-cunha et al., 2015).

In addition to its nuclear DNA, *T. cruzi* has an extranuclear DNA network located in its single mitochondria, called kinetoplast DNA (kDNA) (Rassi et al. al., 2012). This DNA can represent up to 20-25% of the total cellular DNA. The kDNA maxicircles are equivalent to the mitochondrial genome of other eukaryotes and contain the genes coding for rRNA and mitochondrial proteins involved in the electron transport chain (Simpson et al., 1987; Westenberger et al., 2006). Despite the relatively small total length of the molecule (about 40 Kbp), the assembly of this molecule has been difficult because more than half of the DNA sequence is composed of two complex repetitive structures (Gerasimov et al., 2020). The molecule is structured in at least three main compartments, a gene-rich conserved region of about 15 Kbp, a short repetitive region called P5, and a long repetitive region called P12 (Berná et al., 2021). Unit length, number of repeats, and the composition of the repeat sequence differ between the two regions.

*T. cruzi* reference genomes have been highly fragmented and underrepresented for many years due to their genomic complexity and the nature of the data produced by short-read sequencing technologies (El-sayed, et al., 2005; Franzén et al., 2011; Grisard, et al., 2014). For this reason, interest has grown in using long-read technologies to improve the assembly of repetitive regions. In recent years, some *T. cruzi* genomes have been published with these technologies (Supplementary Table 1), improving the understanding of the genome but have also shown the wide variability within and between DTUs and strains of the parasite (Berná et al., 2018; Callejas-Hernández, Rastrojo, Poveda, Gironès, & Fresno, 2018; Wang et al., 2020).

Although knowledge of the *T. cruzi* genome has been expanded in Colombia, there is no detailed description of the composition of the genome that includes the distribution of multigenic families and genetic diversity based on a genome sequenced with long reads technologies. Neither has the complete organization of the kDNA maxicircle been described in Colombian isolates belonging to the DTU TcI. In this paper, the sequencing of *T. cruzi* (TcI) isolated from Colombia was carried out using the new PacBio methodology – High Fidelity (HiFi), and a description of the genome is reported that includes assembly, annotation, and also the genetic diversity between DTUs and comparative genomics.

## RESULTS

### A phased genome assembly for the *T. cruzi* strain Dm25

We performed long read whole genome sequencing of the *T. cruzi* strain Dm25 in exponential growth phase following the PacBio HiFi sequencing protocol. An initial molecular characterization of the strain determined that it belongs to the *T. cruzi* DTU TcI without presenting mixed infection with *T. rangeli* or another *T. cruzi* DTUs. (Supplementary Figure 1). This platform provided 206,520 sequences with an average length of 20,997 bp. The median sequence quality was Q26, with all sequences having a quality greater than Q20. After evaluation of several options for genome assembly, we obtained partially phased assemblies from these reads running the tools Hifiasm and NGSEP. After evaluation and manual selection of the contigs, we built a combined phased assembly for Dm25.

The first haplotype (H1) can be considered a primary assembly and it is composed of 35 contigs representing 32 pseudo chromosomes and one copy of the maxicircle. Almost all chromosomes (30) were assembled in one single contig, and telomeric repeats [(TTAGGG)n] were identified on both chromosome ends for 24 single chromosome contigs and for the two chromosomes assembled in two contigs. The total length of this haplotype is 38.68 Mbp, and the median (N50) length is 1.23 Mbp (Supplementary Figure 2). Performing quality assessment through mapping of conserved genes, only one of the 130 conserved genes in Euglenozoa was fragmented. The remaining 129 genes were uniquely mapped to the assembly.

To assess the ploidy of each chromosome, we realigned the reads to the H1 assembly and we calculated the average read depth for each chromosome, and the distribution of allele dosages for sites having more than one allele call. Figure 1A shows that the average read depth varies around 100x for most chromosomes. Chromosomes 6, 12, 30 and 31 are the main outliers with average read depths ranging from 144x to 192x, suggesting that chromosomes 6, 12 and 30 are triploid and chromosome 31 is tetraploid. Conversely, the average read depth of chromosome 23 is about 51x, which suggests that only one copy of this chromosome is present in Dm25. Based on the average read depth, we predicted that between four and five copies of the maxicircle were present in Dm25. Figure 1B shows the genome wide distribution of relative allele dosages in sites with more than one allele call.

**Figure 1.**
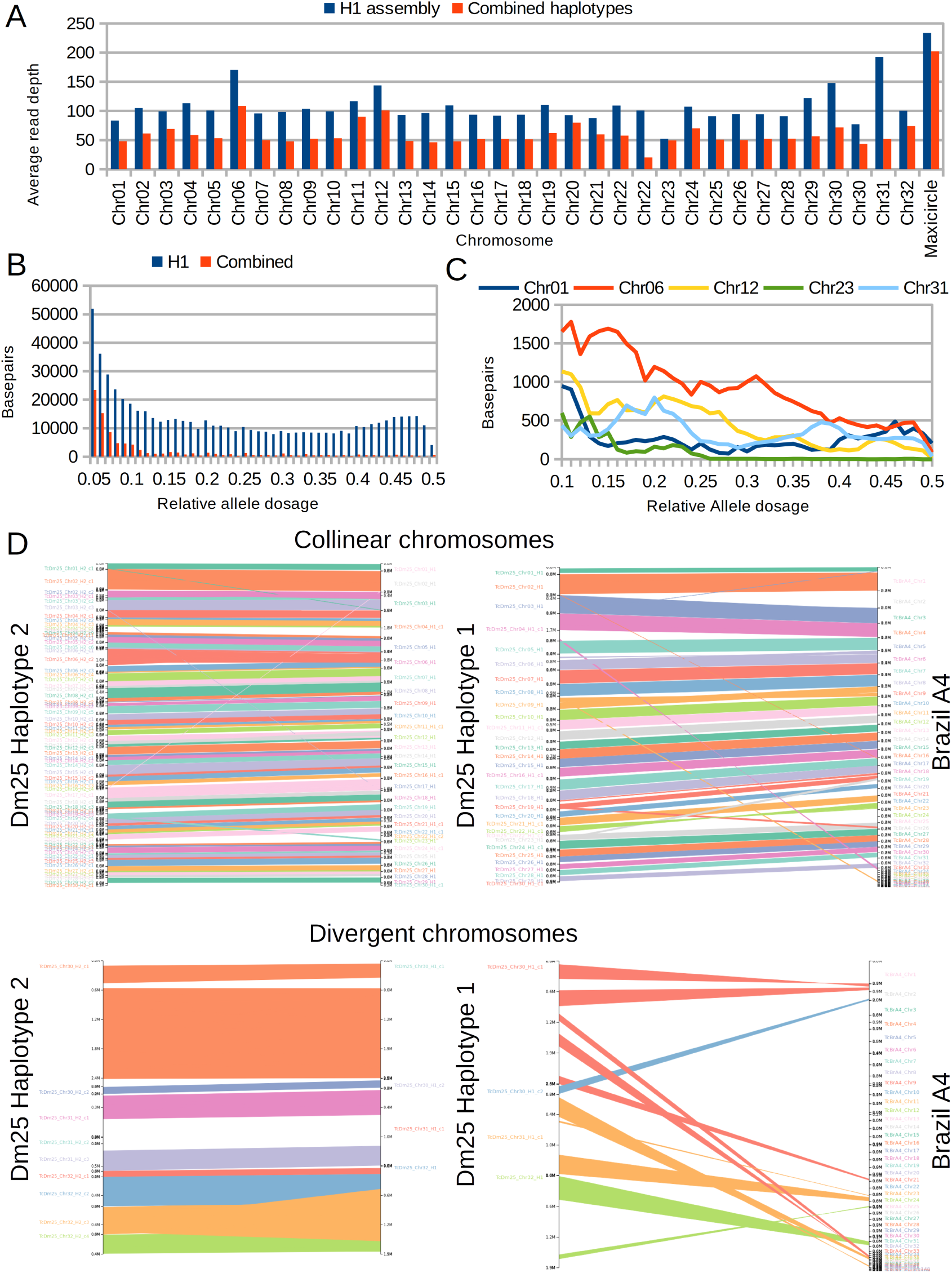
**A**. Average read depth per chromosome of HiFi reads aligned to the primary haplotype and to the complete phased assembly. **B-C**. Histograms of overall allele dosages in sites with two observed alleles. **B.** Complete genome **C.** Individual chromosomes. **D.** Synteny-based alignment between the assembled haplotypes of the genome of Dm25 and between the first haplotype and the Brazil A4 genome assembly.

The peak close to 0.5 suggests that most of the genome is diploid. However, chromosome-specific distributions seem to support the aneuploidies predicted by the read depth distribution. Figure 1C shows these distributions for the presumably aneuploid chromosomes 6, 12, and 31. The figure also shows the distribution for a control diploid chromosome (chr1) and for the haploid chromosome 23. The peak close to 0.33 for chromosome 6 supports that this chromosome has three copies. The peaks close to 0.25 and 0.5 suggest that chromosome 31 is tetraploid. These predictions are consistent with the average read depth. The signal of chromosome 12 suggests a tetraploid chromosome, but the read depth suggests that only three copies are present. A possible explanation for this pattern is that four copies are present for about half of the chromosome.

Considering the expected ploidy for each chromosome, we selected contigs from the automated assemblies to generate a second haplotype (H2). This haplotype was more fragmented having 96 contigs, which represents an average of about three contigs per chromosome. The total length in this case was 37.3 Mbp and the N50 was 572 Kbp (Figure 1D). Only five of the 130 conserved genes in Euglenozoa were not found in the second haplotype. Four of these genes are located in the haploid chromosome 23, which was included only in the first haplotype. The same gene fragmented in H1 was fragmented in this haplotype. Two copies were identified for two of the 124 remaining genes. Taking into account the possible aneuploidies and copy number variation suggested by the read depth analysis, we further generated a partial third haplotype (H3) composed of 83 small contigs adding to 8.05 Mbp. Most sequences (53 sequences adding to 5.32 Mbp) in this haplotype were assigned to the repetitive or polyploid chromosomes 4, 6, 12, 30 and 31. The final genome is a concatenation of these three haplotype assemblies and aims to represent the complete haplotype diversity within the Dm25 genome.

We compared the assembled haplotypes of Dm25 with the genome assembly of the Brazil A4 isolate (Wang et al. 2020) which is the current most accurate haploid assembly of a TcI strain. We did not use this genome to merge contigs to avoid misassemblies produced by true structural variation between Dm25 and A4. However, we used the A4 genome as a reference to sort and orient the chromosomes of the first haplotype that were clearly collinear with A4 chromosomes. We identified two types of chromosomes based on this comparison (Figure 1D). One to one relationships could be identified between chromosomes of Dm25 and chromosomes of A4 for 29 of the 32 chromosomes. The remaining three chromosomes showed a high level of structural variation between the Dm25 and the A4 genome. Chromosome 30 of Dm25 could be mapped to chromosome 2 of A4. However, at least two different syntenies could be identified, suggesting a translocation of this chromosome between the strains. Moreover, this chromosome included two complete copies of the sequences reported as chromosome 21 and chromosome 36 in the A4 genome. Consistent with Wang et al. (2020), we found that this chromosome is almost entirely composed of copies of the most repetitive gene families in *T. cruzi*. Chromosome 31 of Dm25, could be mapped partially to chromosome 24 of A4. However, it also included complete copies of the sequences labeled as chromosome 37 and chromosome 43 of A4. The two chromosome copies of Dm25 could be assembled in one and three contigs respectively. However, a large divergence was observed between the two copies in the central part of the chromosome. Finally, chromosome 32 of Dm25 (haplotypes assembled in one and four contigs) included segments syntenic with sequences termed as chromosomes 25, 32, 35, 38 and 39 of A4.

### Gene annotation and protein evolution in *Trypanosoma* species

We identified 47,772 transposable elements (TEs) in the combined assembly, covering 36.4 Mbp (44%) of the assembly (Supplementary Table 2). Figure 2A displays the density of repetitive elements across the H1 haplotype. The percentage of the genome covered by TEs was larger for the haplotype H1 (48%), compared to H2 (38%) and H3 (35%), probably because a larger number of copies of TEs could be assembled in the contigs included in H1. Most TEs (25,355 covering 24.3 Mbp of the genome) were annotated as “Unknown”.

**Figure 2.**
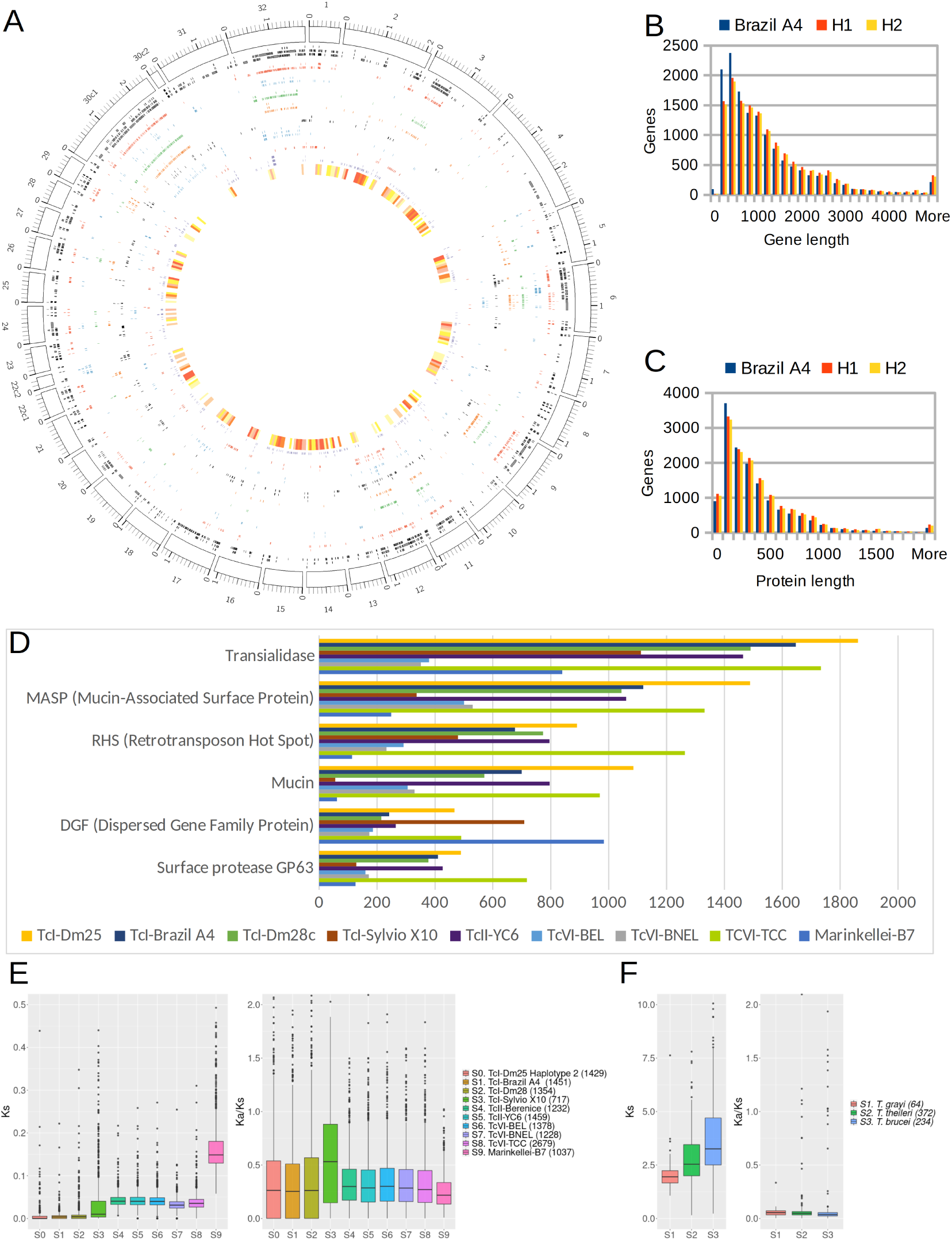
**A.** Distribution of genes and repeat elements in the primary assembly of the Dm25 strain. Tracks show repeat density, Transialidases, Transposable element domains, Mucin-Associated surface proteins, Mucines, Retrotransposon hot Spots, Surface proteases, Dispersed gene families, Kinases and density of single copy genes. **B-C**. Length distribution for genes and proteins annotated in the two haplotypes of TcDm25, and in the Brazil A4 strain. **D.** Number of copies of the six main gene families, compared with those reported in previous studies. **E-F**. Distributions of synonymous nucleotide divergence rate (ks) and relative non-synonymous to synonymous divergence rate (ka/ks) for pairs of orthologs, comparing the primary assembly of TcDm25 (H1) with other *T. cruzi* assemblies (**E**) and with assemblies of other species (**F**).

Following the pipeline implemented in the Companion website (Steinbiss et al., 2016), we annotated 29,544 genes for the complete assembly. The completeness of H1 translated into a larger number of annotated genes (14,207), compared to H2 (13,679). The gene catalog was complemented with 1,658 additional genes annotated in H3. Compared to the available annotation of the Brazil A4 strain, the annotation of each haplotype has about the same number of genes. However, the lengths of the genes annotated in H1 and H2 (about 1.4Kbp) are on average about 200bp larger than the lengths of the genes annotated in A4 (Figure 2B). Consequently, proteins annotated in the two haplotypes are about 45 amino acids longer than those annotated in A4 (Figure 2C).

Combining protein family annotations with homologies between Dm25 and annotated genes in the A4 transcriptome, we found a large and consistent number of copies of the main protein families: Transialidase (TS), Mucin-Associated Surface Protein (MASP), Retrotransposon Hot Spot (RHS), Mucin, Dispersed Gene Family (DGF) and Surface Protease GP63. Figures 2A and 2D show the distribution of these gene families across the genome and the number of genes per family. Based on protein domains and orthology, we also identified large families of protein kinases (PKS) and core histone proteins (HIS). We observed a large overlap between the TS, MASP, RHS, DGF, and MUC families with annotated TEs. In contrast, the protein kinases did not overlap with annotations of transposable elements.

We analyzed the nucleotide and protein evolution between genes annotated in our assembly and genes annotated in other *T. cruzi* assemblies. Figure 2D shows the normalized differences in synonymous sites (Ks) comparing synteny orthologs of the haplotype H1 with other *T. cruzi* assemblies, including the haplotype H2. The Ks values are on average below 0.02 for comparisons with TcI assemblies, although a larger variance is observed in the comparison with the strain Sylvio, compared to H2, A4 and Dm28. The Ks values obtained from orthologs with assemblies of other DTUs are significantly higher than those obtained from comparisons within TcI (p-value < 10^-16^ of a Wilcoxon rank test). However, the absolute values are on average lower than 0.05, except for the comparison against the assembly of a *T. cruzi* marinkellei strain, for which the average increases to 0.15. In contrast, the Ks values of synteny orthologs between the first haplotype of Dm25 and the genes annotated in other species are on average above 2 (Figure 2E). This suggests high divergence times between *T. cruzi* and other species with contiguous assemblies (*T. grayi*, *T. thelleri* and *T. brucei*).

We also investigated the behavior of protein evolution for single copy synteny orthologs within and between species. For each pair of unique synteny orthologs, we calculated the ratio between normalized non-synonymous and synonymous mutations (Ka/Ks). According to the neutral theory of evolution, this value should be close to 1 for genes not affected by selection. Values lower than 1 indicate purifying selection and values higher than 1 indicate positive selection and rapid protein evolution. Most Ka/Ks values for orthologs within species fall below 1 with averages between 0.3 and 0.5 (Figure 2D), which suggests that most core genes are subject to some level of protein conservation through purifying selection. Outliers have Ka/Ks>1 and are genes with rapid protein evolution. Conversely, the Ka/Ks values for orthologs between species were on average lower than 0.1 (Figure 2E). This result which at first sight looks surprising can be explained because very few orthologs could be identified between these species, and hence these genes are probably those with the highest selective pressure for protein conservation.

Regarding multicopy gene families, Figure 3A shows the distribution of Ks and Ka/Ks values, differentiating paralogs by physical proximity in the genome. The Ks values were significantly lower for close paralogs, compared to distant paralogs, suggesting that tandem duplications are more recent than interspersed duplications. The difference was larger for Mucin and especially for DGF paralogs, compared to other families. This suggests that the diversification and spread of the DGF paralogs across the genome happened at a much older time compared to other families. Ka/Ks values were closer to 1 compared to the values obtained from single copy orthologs, suggesting that these genes are subject to a more relaxed purifying selection, compared to single copy genes. The averages for both close and distant paralogs of the MASP and Mucin families were larger than 1, suggesting a faster protein evolution, compared to the other families. Conversely, close paralogs of the TED, PKS and HIS families and distant paralogs of the DGF and HIS families have a Ka/Ks distribution similar to that observed for single copy orthologs. The latter case suggests that purifying selection is acting to preserve the protein sequence of DGF and HIS paralogs.

**Figure 3.**
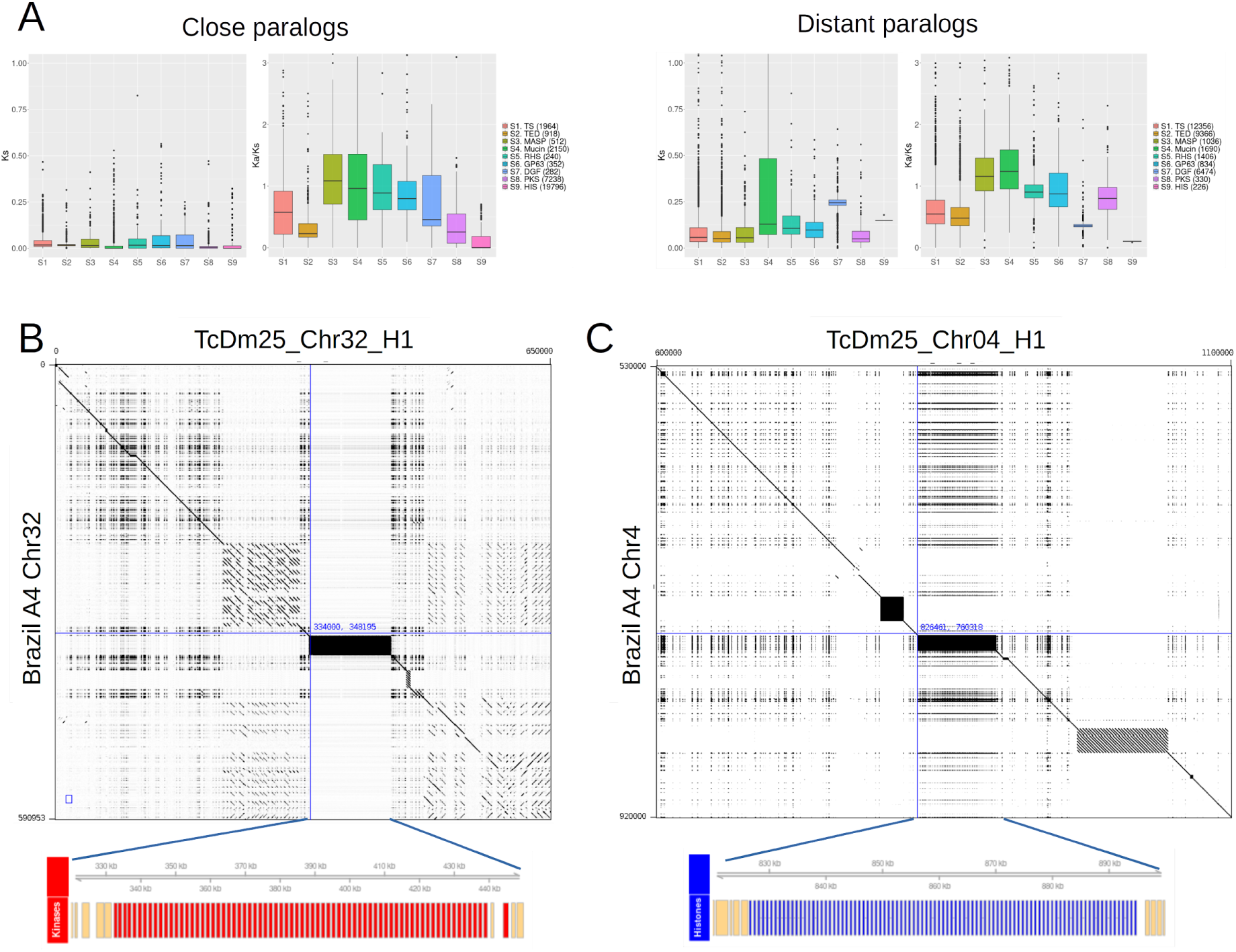
**A.** Distributions of synonymous nucleotide divergence rate (ks) and relative non-synonymous to synonymous divergence rate (ka/ks) for pairs of paralogs of the six main multicopy gene families, and for protein kinases (PKS) and histones (HIS), discriminating tandem (close) paralogs from distant paralogs. **B**. Alignment of chromosome 32 of the haplotype H1 of Dm25 with chromosome 32 of Brazil A4. The lower track highlights a tandem array of protein kinases spanning the black rectangle. **C.** Alignment of chromosome 4 of the haplotype H1 of Dm25 with chromosome 4 of Brazil A4. The lower track highlights a tandem array of histones spanning the black rectangle.

In contrast to the gene families described in previous studies, close paralog pairs of protein kinases (7,238) were much abundant than distant paralog pairs (330). The reason for this is that most genes in this family resulted from a recent tandem duplication generating 71 copies of protein kinases spanning 200 kbp of chromosome 32 (Figure 3B). The syntenic region in the haplotype H2 contains 67 homolog copies (Supplementary Figure 3). This duplication event was not clearly observed in previous genome assemblies. A tandem array of 9 homologs in the Brazil A4 assembly are located in the syntenic region of the contig termed Chr32 (Figure 3B). Besides this, 20 additional homologs are located in three contigs smaller than 32 kbp and not assigned to chromosomes. This suggests that a similar expansion could be present in the Brazil A4 assembly but that it could not be completely reconstructed due to technical limitations of the sequencing technology or the assembly pipeline. The comparison with the genome assembly of the Dm28 strain revealed a similar situation (Supplementary Table 3). Tandem arrays of 28 and 8 copies were identified in two contigs of 101 kbp and 22 kbp respectively. Conversely, no orthologs were identified in the Sylvio genome assembly. Comparing with assemblies of strains belonging to other DTUs, tandem arrays of 12, 22, and 16 homologs were observed in one contig of the Berenice assembly and two contigs of the TCC assembly.

The annotation also revealed two large recent tandem duplications of core histones, located in chromosomes Chr04 (69 copies in H1, 33 copies in H2, Supplementary Figure 4) and Chr18 (46 copies in H1, 43 copies in H2). This expansion is not observed in the Brazil A4 assembly (Figure 3C). The assembly of the Sylvio strain was the only TcI assembly in which homolog tandem arrays were identified, having 95 copies homolog to the array on Chr04 and 34 copies homolog to the array in Chr18 (Supplementary Table 3). Two tandem arrays of 21 and 44 genes, both homologous to the array located on Chr18 were identified in two separate contigs of Dm28. Regarding other DTUs, only the TCC assembly has three tandem arrays of 42, 19 and 22 genes. While the first array is homologous to the array on Chr04 of Dm25, the other two arrays are homologous to the array on Chr18.

### Complete reconstruction of the *T. cruzi* (TcI) Dm 25 maxicircle

Kinetoplastid molecules were sampled in the genome assembly of Dm25, obtaining a complete reconstruction of the maxicircle. After circularization and sorting, the total length of this assembly was 47,166 bp (Figure 4). Performing gene annotation we identified 18 protein coding genes: the subunits ND1, ND3, ND4, ND5, ND7, ND8, ND9 of the NADH dehydrogenase, the cytochrome B (CyB), the subunits I, II y III of the cytochrome c oxidase (COI, COII, COIII), the ATPase six (ATP6), the ribosomal protein S12 (RSP12), and five genes with unknown function (MURF1, MURF2, MURF5, CR3, CR4). We also found two ribosomal RNA genes (12S, 9S). The 12S gene was used to mark the start of the circular sequence in the assembly. These genes make up the entire conserved region of the Maxicircle, which is consistent with the maxicircle structure defined by Ruvalcaba-Trejo & Sturm (2011). Gene lengths and orientations were similar to those reported in previous studies (Callejas-Hernández et al., 2021; Westenberger et al., 2006; Ruvalcaba-Trejo & Sturm, 2011; Berná et al., 2021). The total length of the conserved region is 15,429 bp (32.7% of the total) and the GC-content is 25.34%. Most of the region has an excess of cytosine relative to guanine in the positive strand, which is measured by a negative GC-skew statistic (Westenberger et al., 2006). The seven genes that are annotated in the negative strand (ND9, MURF5, MURF1, ND1, COI, CR4 and ND3) overlap with segments showing a neutral or a positive GC-skew.

**Figure 4.**
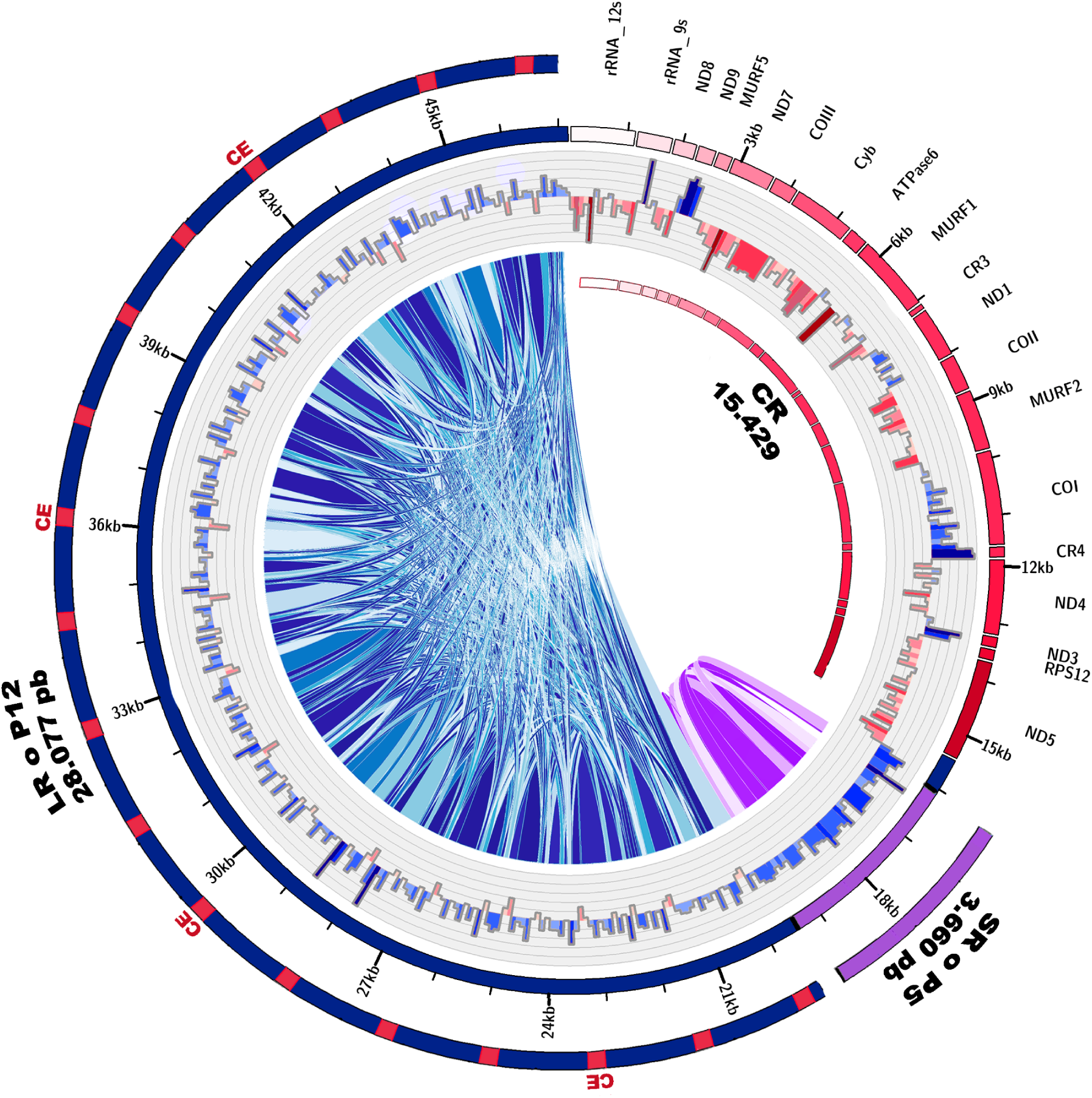
Assembly of the maxicircle of the *T. cruzi* strain Dm25. Protein coding genes in the conserved region (CR) are shown in the external red rectangles. The divergent region is divided into the P5 region (purple) and the P12 region (blue). Conserved elements (CE) across the P12 region are shown as red rectangles. The central histogram shows the GC-skew, positive values are in red and negative values are shown in blue (windows size 100 bp). Internal bands show the homology relationships making up the P5 and P12 variable regions.

The remaining 31,737 bp (67.3% of the assembly) corresponds to the variable region. The GC-content of this region (21.39%) is lower than that of the conserved region. Following previous works (Berná et al. 2021, Gerasimov et al., 2020), we divided this region into a small subregion (3,660 bp), termed P5, and a long subregion (28,077 bp), termed P12. The two regions are composed of a series of tandem repeats with unit lengths around 150bp for the short region, and 1,500bp for the long region. Nucleotide identity between pairs of repeat copies ranged from 75.6% to 100%. Comparing the P12 region with previous assemblies, we identified 18 palindromic conserved elements, which are known to be involved in different molecular processes such as the replication of the mitochondrial DNA (Gerasimov et al., 2020).

### Genetic variability in *T. cruzi*

To evaluate the use of our haploid genome assembly as a reference for diversity studies, we reanalyzed publicly available WGS Illumina reads sequenced from 39 *T. cruzi* strains classified in different DTUs (Supplementary Table 4). The number of reads of the downloaded sequences was highly variable and the mapping rate ranged from 34% (2 samples) to 86% (Supplementary Figure 5).

A raw dataset of 1,018,520 single nucleotide variants was found in all Illumina sequences mapped to the first haplotype of the Dm25 assembly. Figure 5A shows that the TcI group has the least number of variants, except for the H1 strain from Panama, which has about 6 times more variants. The number of variants of the H1 strain is similar to the variants of the TcV group. Strains Berenice and 9280cl.2 had a significantly lower number of variants compared to their groups, this is due to the low number of reads sequenced in these strains (Supplementary Figure 4). In general, these results are consistent with previous molecular characterization showing that the Dm25 strain belongs to the TcI group.

**Figure 5.**
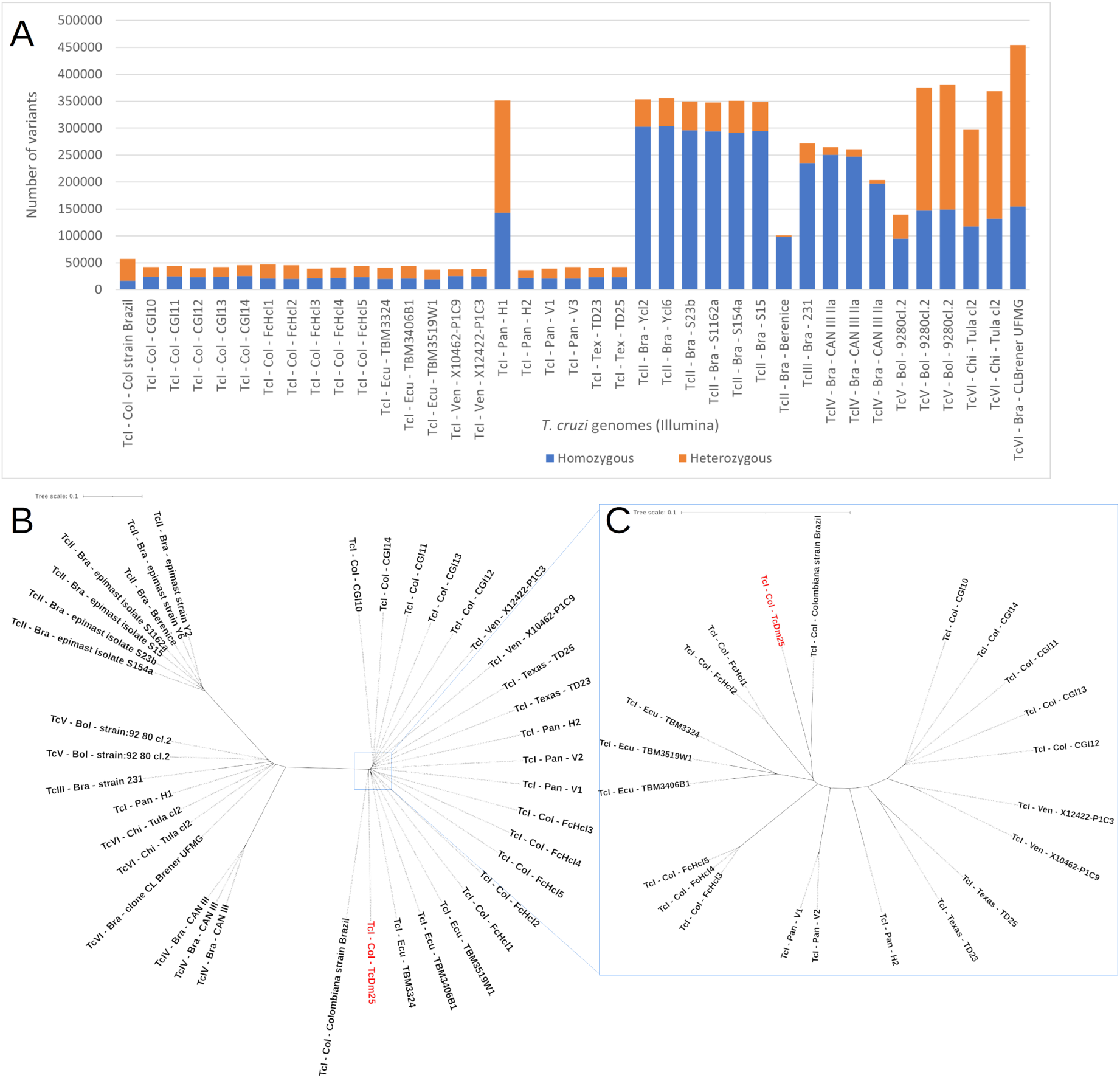
Genomic variants identified between Illumina reads of *T. cruzi* strains from different DTUs and our haploid *T. cruzi* assembly (H1). **A.** Number of homozygous and heterozygous variants in *T. cruzi* strains. **B.** Neighbor joining clustering of genetic distances between *T. cruzi* strains from different DTUs, including the haploid TcDm25 assembly. **C.** close up of the variability within TcI. The names indicate the DTU – country of isolation – strain. Bra: Brazil, Bol: Bolivia, Pan: Panama, Chi: Chile, Col: Colombia, Ven: Venezuela, Ecu: Ecuador.

We derived a neighbor joining clustering from the variants identified in the *T. cruzi* strains with Illumina reads (Figure 5B). All the strains from TcI were grouped in the same node, except for the H1 strain from Panama which is consistent with Figure 5A. Likewise, the TcII strains were grouped on the left side of the clustering. The TcIII, TcV, and TcVI groups were grouped in a node, together with the TcI strain from Panama. This node looks intermediate between TcI and TcII groups. Although the reads of Dm25 were not obtained by Illumina technology, we were able to confirm that Dm25 strain belongs to the TcI group with this analysis.

Figure 5C shows a close look at the clustering within the TcI group. The CGl strains, which are parasites isolated from a patient with HIV and cardiomyopathy, are clustered together on the right side of the figure. The FcHcI strains, which correspond to parasites isolated from an acute chagasic patient infected by oral transmission, formed two distinct groups. The Colombiana strain was placed close to Dm25 but the separation suggests a level of divergence between these strains larger than that observed within the other groups. In general, a grouping by countries is observed, except for the strains from Colombia.

## DISCUSSION

The development of sequencing methods to obtain high-quality long read DNA sequencing data enabled the complete characterization of complex genomes, including phase reconstructions for diploid and even polyploid species. In this work, we report the nearly complete and phased assembly of a Colombian TcI strain of *T. cruzi*. The use of the PacBio HiFi technology allowed us to annotate and analyze a complete catalog of the most important repetitive structures, including transposable elements and multicopy gene families. Compared to previous efforts using the Oxford Nanopore technology (Wang et al., 2021), the small error rate of HiFi reads allowed us to identify heterozygous sites, make inferences about ploidy per chromosome, and reconstruct two haplotypes for diploid and aneuploid chromosomes. A wide variability between the haplotypes of TcI and even within the TcI haplotypes was evidenced, especially for three chromosomes, which are enriched for multicopy gene families. This confirms the broad plasticity of the genome and the strain-specific evolution of *T. cruzi* previously reported (Berná et al., 2018; Callejas-Hernandez et al., 2018; Díaz-Viraqué et al., 2019; Talavera-Lopez et al., 2021; Wang et al., 2021). Further improvements in read quality and length of long read sequencing technologies will facilitate the full reconstruction of large numbers of strains of genomes for pathogens with complex genomes such as those in the *Trypanosomatidae* family. This will allow researchers to characterize and analyze the function and evolution of the expectedly large haplotype variability segregating within *T. cruzi*.

The total length of the genome assembly of Dm25 (84 Mbp) is consistent with previous flow cytometry experiments in which the total genome size was estimated to fluctuate between 80 and 150 Mbp (Lewis et al., 2009). The percentage of the genome covered by repetitive elements (∼47%) is also consistent with previous reports (Reis-Cunha et al., 2015: Wang et al., 2021), and only differs from the percentage reported for the Sylvio X10 strain, which was only 18.43% (Talavera-Lopez et al., 2021). Even using long reads, heterozygosis and a large percentage of repetitive elements are the main difficulties to obtain chromosome-level genome assemblies (Jarvis et al., 2022). Part of the complexity of the *T. cruzi* genome is evidenced by the presence of different aneuploidies. Recent studies also support the presence of aneuploidies in the genomes of TcI strains (Cruz-Saavedra et al., 2022). In particular, chromosome 31 seems to have a consistent increase in number of copies, compared to diploid chromosomes (Reis-Cunha et al., 2015). One of the characteristics of this chromosome is the abundance of genes related to protein glycosylation, such as mucin surface proteins. These proteins can be related to the survival of *T. cruzi* during the infection process (Buscaglia et al., 2006; De Pablos et al., 2012). Previous studies also have investigated the role of aneuploidies to facilitate a rapid adaptation of the pathogen across its life cycle, while moving from an invertebrate to a vertebrate host, through modulation of allele dosages (Dujardin et al., 2014; Reis-Cunha et al., 2018).

The number of gene copies per family identified within each haplotype was consistent with previous studies (Berná et al., 2018, Wang et al., 2021). The analysis of nucleotide and protein evolution over paralog pairs provides insights into the relative times of expansion of the different families, and their level of protein conservation. With the exception of Mucins and dispersed gene families (DGF), the ks value in most comparisons between paralogs was below 0.2. The similarity of these distributions with the distribution of ks values for core orthologs against the *T. cruzi* marinkellei assembly, suggests that most of the observable expansions of gene families occurred along the diversification of the species. As expected, tandem paralogs seem to have appeared at a smaller time, compared to distant paralogs. The distribution of ka/ks values suggests that protein sequences of multicopy gene families evolve faster than core genes, thanks to a more relaxed purifying selection. A notable exception to this pattern is the case of distant copies of dispersed gene families (DGF). The expansion of this family seems to occur at a much older time, compared to the other families, but at the same time, the ka/ks values suggest a high level of protein conservation. High protein conservation and lack of positive selection in this family were previously reported in a study in which Shannon entropy was used as a measure of variability across a protein sequence alignment (Kawashita et al., 2009). This is surprising taking into account that subtelomeric regions contain copies of DGF genes (Moraes et al., 2012). These regions usually have higher levels of homologous recombination and are even subject to ectopic recombination, which increases the variability of genes present in these regions (Christiaens et al., 2012). Moreover, recent studies reported that many subtelomeric coding sequences of DGF genes serve as replication origins (de Araujo et al., 2020), and that up to 80% of DGF genes include dynamic nucleosomes (Lima et al., 2021). It has also been shown that DGF genes are expressed in the three life cycle stages (Kawashita et al., 2009, Lander et al., 2010). Further studies are needed to elucidate why the pathogen requires protein conservation for this family.

Beyond the well characterized repeat families, we observed three recent expansions of protein kinases and histones. A comparative analysis of public assemblies within the syntenic regions that could be identified suggests that these expansions are not a unique feature of TcDm25, but that the expansions could not be fully characterized due to the lack of completeness of previous assemblies. Gene copies of the expansion of protein kinases in chromosome 32 belong to the TcCK1.2 gene family. This is a casein kinase 1 (CK1) which is a signaling serine/threonine protein. These proteins are involved in different cellular processes such as protein trafficking, cell cycle regulation, cytokinesis, DNA repair and apoptosis (Spadafora et al., 2002; Knippschild et al., 2005). TcCK1.2 is more expressed in the amastigote stage, compared to the epimastigote and the trypomastigote stages (Spadafora et al., 2002). Orthologs TcCK1.2 in *L. donovani* (Rachidi et al., 2014) and *T. brucei* (Urbaniak., 2009) are crucial for parasite survival in the amastigote stage. This suggests that the expansion of TcCK1.2 could be related to an adaptation mechanism. Likewise, gene copies of the expansion of histones found in chromosome 4 belong to the histone variant H2B.V. Histones play a key role in the organization of the chromatin structure and gene expression in *T. cruzi* (de Lima et al., 2020). Previous studies analyzing chromatin extracts (de Jesus et al., 2017), ChIP-seq data and performing knockout experiments (Roson et al., 2022) showed that this variant is associated with nucleosome instability and that it is more expressed in epimastigotes compared to trypomastigotes. This suggests that H2B.V genes can be related to chromatin structure changes and modulation of transcription rates (Elias et al., 2001). Further functional experiments are needed to reveal the relationship between gene copy number and expression changes during host-pathogen interactions.

Nucleotide evolution statistics on core orthologs suggest that there is a large gap in the range of species that need to be sequenced to obtain a full reconstruction of the evolutionary history of *Trypanosomatidae*. The closest species to *T. cruzi* that we could identify with a publicly available genome was *T. grayi*. The average ks values for core ortholog pairs were close to two, suggesting a very large divergence time between these species. High protein conservation was observed in these paralogs, suggesting that only ultraconserved essential proteins were included in this comparison.

The use of high-quality long reads allowed to obtain a complete and direct assembly of the maxicircle, without any scaffolding or curation steps. The complex pattern of long and short repetitive elements present in the divergent region explains why the Maxicircle can not be fully reconstructed using short read technologies (Urrea et al., 2019; Lin et al., 2015). The total length of our assembly (47 Kbp) is within the range between 35 Kbp and 51 Kbp estimated by previous studies (Berná et al., 2021). The organization of the molecule in a gene rich conserved region and two divergent and repeat rich regions is also consistent with previous assemblies. Previous studies in *T. brucei* showed that this region contains binding sites for the topoisomerase II, which indicates that this region is important for the replication of the molecule (Myler et al., 1993) . Recent studies in *T. vivax* indicate that the variable region can have a large variability within species because copies of repetitive elements can recombine, producing presence/absence variants and even rearrangements (Greif et al., 2021)

Based on the reanalysis of publicly available Illumina data, we show that the primary assembly of TcDm25 can serve as a reference for population genomic studies in *T. cruzi*. A wide variation in the number of variants in the different DTUs was evidenced, which is consistent with the reported genomic heterogeneity (Talavera-Lopez et al., 2021; Wang et al., 2021). Sample clustering based on SNP variation separates the reported DTUs (Zingales et al., 2009). A single NJ cluster includes the TcIII, TcV, and TcVI groups because TcV and TcVI are hybrids of TcIII and TcII (Zingales et al., 2009) although a more comprehensive sampling within DTUs is needed to corroborate this hypothesis. The genetic proximity between TcIII, TcV and TcVI strains has been reported in previous studies with markers such as gGAPDH where it has been impossible to separate hybrid strains from parental strains (Brandão et al., 2020). In addition, it was possible to corroborate the erroneous assignment of the H1 strain (from Panama) to the TcI group (Majeau et al., 2021). We expect that this resource will be very valuable for different research groups performing evolutionary, functional and population genomic analysis in *T. cruzi* and other related tropical pathogens.

## METHODS

### Sampling area and parasite culture

The capture of the host *Didelphis marsupialis* was carried out in the municipality of Coyaima, department of Tolima (coordinates 3.8025-75.19833), using baited tomahawk traps (Supplementary Figure 6). The collected specimen was sedated and individualized, to later take a blood sample, with the purpose of performing a blood culture in a biphasic medium (NNN: Novi, Nicolle, McNeal / LIT: Liver Infusion Tryptose). The strain of *T. cruzi* isolated was then cryopreserved in liquid nitrogen at the Laboratorio de Investigaciones de Parasitología Tropical (LIPT) of Universidad del Tolima until use. The isolate identified as Dm25 was thawed and placed in NNN culture medium with LIT supplemented with 15% fetal bovine serum (FBS) and 100 IU/ml gentamicin/ampicillin mixture at 28° C for its log-phase growth, allowed obtaining 10^8^parasites for the molecular characterization and whole genome sequencing (WGS).

### Molecular characterization

Species identification was based on the amplification of the hypervariable region of trypanosomatid minicircles using primers S35 (5-AAA TAA TGT ACG GGT GGA GAT GCA TGA-3), S36 (5-GGG TTC GAT TGG GGT TGG TGT-3) and KP1L (5-ATA CAA CAC TCT CTA TAT CAG G-3) as proposed by Vallejo et al. (2002). Finally, the amplification products were visualized by electrophoresis with 6% polyacrylamide gels, stained with silver nitrate, and 1kb Plus DNA Ladder (Invitrogen ™ by ThermoFisher Scientific, Product 10787018).

For the genotyping of the *T. cruzi* isolate within the Discrete Taxonomic Unit corresponding to lineage I (DTU I) or II (DTU II), the intergenic region of the spliced-leader gene (SL-IR) was amplified using the primers proposed by Souto et al. (1996), TCC/TCI/TCII, which amplifies a product of 300 bp corresponding to DTU II and 350 bp corresponding to DTU I. The amplification products were visualized in agarose gel electrophoresis stained with 2% Ethidium Bromide.

### DNA extraction and genome sequencing

DNA extraction from *T. cruzi* epimastigotes was performed using the Gentra Puregene kit (Qiagen) to obtain high molecular weight DNA. The DNA was quantified with the NanoDrop 2000 spectrophotometer (Thermo Scientific, USA). The integrity of the DNA was verified by electrophoresis with a 2% agarose gel (90V for 30 minutes). *T. cruzi* DNA sequencing was performed using Pacific Bioscience (PacBio) HiFi technology at 100X average read depth.

### Sequence assembly and quality assessment

An initial phased *de novo* assembly of the *T. cruzi* strain Dm25 was performed running Hifiasm v0.12(r304) (Cheng et al., 2021) and the Next Generation Sequencing Experience Platform (NGSEP) v4.3.1 using the Assembler command with k-mer length 25, window length 40 and ploidy of 2 (Gonzalez-Garcia et al. 2023). Contigs were aligned to the publicly available haploid genome assembly of the TcI Brazil A4 strain using Minimap2 v2.22 (Li et al., 2018). Based on these alignments, the contigs of both haplotypes were manually sorted and for each chromosome contigs making the haplotypes H1 and H2 were selected manually. We aligned the PacBio reads to a concatenated H1/H2 assembly, called heterozygous variants using NGSEP, and calculated mean read depths using samtools v1.16 (Danecek et al., 2021) to determine genomic regions showing evidence of the presence of a third haplotype. Unassigned contigs were also mapped to the H1/H2 assembly to select contigs likely to belong to the third haplotype. Reads were mapped again to a concatenated genome including the H3 contigs to validate that the H3 contigs will have good read coverage.

Genome statistics were obtained running QUAST v5.02 (Gurevich et al., 2013). Per base quality assessment through mapping of conserved genes was assessed using BUSCO v5.3.2 searching the reference dataset of 130 genes in Euglenozoa (Manni et al., 2021).

### Genome Annotation

We generated an initial database of Transposable elements using Repeat Modeler (Flynn et al., 2020). We mapped this initial database to the genome using Repeat Masker (http://www.repeatmasker.org/). We also executed the TransposonsFinder command of NGSEP (Gonzalez-Garcia et al., 2023b) to map the database generated with Repeat Modeler and annotate the regions including transposable elements in the genome assembly of *T. cruzi* (TcDm25). We executed a separate process for each haplotype and executed two rounds of identification.

We ran the Companion software for structural and functional annotation of the genome of *T. cruzi* (Steinbiss et al., 2016). Companion combines the tool RATT (Otto et al., 2011) to transfer models of publicly available assemblies, with ab-initio predictions obtained with SNAP (Korf, 2004), and AUGUSTUS (Stanke et al., 2006). Functional annotation is performed by transferring annotations from orthologs obtained running OrthoMCL (Li et al., 2003), and performing blast searches to the Pfam-A database (Finn et al., 2014). Additionally, this software takes into account that the genes of kinetoplastids such as *Trypanosoma* and *Leishmania* are organized in large directional groups of genes that are transcribed together as polycistrons. Hence, this software has a filtering method to eliminate the over-prediction of genes on the complementary strand. Genes belonging to multicopy gene families were identified combining genes with direct annotations of characteristic PFam domains with orthologs of genes in the Brazil A4 strain belonging to the family.

### Ploidy determination

Absolute chromosomal ploidy of *T. cruzi* assemblies was determined by estimating allele frequencies from the proportion of occurrence of each heterozygous site using the RelativeAlleleCounts command in NGSEP (Urrea et al., 2018).

### Genome alignments and sequence evolution statistics

Pairwise genome-wide comparisons between the haplotypes of Dm25 and among the publicly available genome assemblies included in this study were performed running the GenomesAligner command of NGSEP (Tello et al., 2023). Assemblies were selected from the TritrypDB database of VEuPathDB (Amos et al., 2021) based on their contiguity, which is related to the use of Sanger or long-read technologies. Dotplots of local alignment were performed using Gepard v1.40 (Krumsiek et al., 2007). To calculate nucleotide and protein evolution statistics (Ks and ka/ks) the DNA coding sequences of homologs inferred from each pairwise comparison were aligned keeping codon information and the command codeml of paml v4.9j (Yang, 2007) was used.

### Determination of variants between DTUs of T. cruzi

To determine variants between *T. cruzi* genomes from different DTUs, 33 Illumina-sequenced *T. cruzi* genomes were obtained from the TriTrypDB database (https://tritrypdb.org) (Supplementary Table 4) (Majeau et al., 2021). The mapping of Illumina reads to TcDm25 H1 haplotype assembly was performed using NGSEP 4.2.1. The same tool was used to call the variants, the functional annotation of the variants, the genotyping quality filter and to obtain the statistics. Finally, a distance matrix was made from the variants and a dendrogram in the iTOL tool (Letunic & Bork, 2021).

### Maxicircle genome assembly and annotation

We used BLAST+ (v2.11.0) (Altschul et al., 1990) to search known maxicircle sequences in the assembly obtained with NGSEP. Known maxicircles were downloaded from the databases NCBI nucleotide and TriTrypDB. We used EMBOSS (Rice et al., 2000) to filter out contigs with GC-content less than expected. The maxicircle was manually annotated, looking for base pair level synteny between the assembly and the annotated maxicircle sequences using BLAST and ARTEMIS v. 18.1.0 (Carver et al., 2012). These tools were also useful to identify and deduplicate repeated extremes and to orient the contig. BLAST+ v2.11.0) was also run with a maximum e-value of 10-6 to find tandem repeats and define the variable regions. The annotated sequence was visualized using Circos v0.69 (Krzywinski et al., 2009).

## DATA AVAILABILITY

The data used in this study is available at the NCBI sequence read archive (SRA) database (https://www.ncbi.nlm.nih.gov/sra) with bioproject accession number PRJNA994590. The genome assembly is available at the Assembly database of NCBI (https://www.ncbi.nlm.nih.gov/assembly/) within the same bioproject accession number.

## ACKNOWLEDGMENT

The work presented in this manuscript was funded by the Colombian Ministry of Sciences research fund “Patrimonio Autónomo Fondo Nacional de Financiamiento Para la Ciencia, la Tecnología Y la Innovación Francisco José de Caldas”, through the grant with contract number 80740-441-2020, awarded to JD. We also acknowledge the IT Services Department and ExaCore of the Vice Presidency for Research & Creation at Universidad de Los Andes for their technical support to perform the computational analysis.

## REFERENCES

Altschul, S. F., Gish, W., Miller, W., Myers, E. W., & Lipman, D. J. (1990). Basic local alignment search tool. Journal of molecular biology, 215(3), 403–410. https://doi.org/10.1016/S0022-2836(05)80360-2

Amos B, Aurrecoechea C, Barba M, et al. (2021). VEuPathDB: the eukaryotic pathogen, vector and host bioinformatics resource center, Nucleic Acids Research 50(D1): D898–D911. https://doi.org/10.1093/nar/gkab929

Barnabé, C., Mobarec, H. I., Jurado, M. R., Cortez, J. A., & Breniére, S. F. (2016). Reconsideration of the seven discrete typing units within the species Trypanosoma cruzi, a new proposal of three reliable mitochondrial clades. Infection, Genetics and Evolution, 39, 176–186. https://doi.org/10.1016/j.meegid.2016.01.029

Berná, L., Rodriguez, M., Chiribao, M. L., Parodi-Talice, A., Pita, S., Rijo, G., … Robello, C. (2018). Expanding an expanded genome: long-read sequencing of Trypanosoma cruzi. Microbial Genomics, 4(5). https://doi.org/10.1099/mgen.0.000177

Berná, L., Greif, G., Pita, S., Faral-Tello, P., Díaz-Viraqué, F., Souza, R. D. C. M. D., Vallejo, G., Alvarez-Valin, F. & Robello, C. (2021). Maxicircle architecture and evolutionary insights into Trypanosoma cruzi complex. PLoS neglected tropical diseases, 15(8), e0009719.

Brandão, E., Xavier, S. C., Rocha, F. L., Lima, C. F., Candeias, Í. Z., Lemos, F. G., … & Roque, A. L. (2020). Wild and Domestic Canids and Their Interactions in the Transmission Cycles of Trypanosoma cruzi and Leishmania spp. in an Area of the Brazilian Cerrado. Pathogens, 9(10), 818.

Buscaglia CA, Campo VA, ACC F, Di Noia JM. (2006). Trypanosoma cruzi Surface mucins: host-dependent coat diversity. Nature Reviews Microbiology 4: 229–236.

Callejas-Hernández, F., Rastrojo, A., Poveda, C., Gironès, N., & Fresno, M. (2018). Genomic assemblies of newly sequenced Trypanosoma cruzi strains reveal new genomic expansion and greater complexity. Scientific Reports, 8(1), 1–13. https://doi.org/10.1038/s41598-018-32877-2

Callejas-Hernández, F., Herreros-Cabello, A., del Moral-Salmoral, J., Fresno, M., & Gironès, N. (2021). The complete mitochondrial DNA of Trypanosoma cruzi: Maxicircles and minicircles. Frontiers in cellular and infection microbiology, 556.

Carver, T., Harris, S. R., Berriman, M., Parkhill, J., & McQuillan, J. A. (2012). Artemis: an integrated platform for visualization and analysis of high-throughput sequence-based experimental data. Bioinformatics, 28(4), 464–469.

Cheng, H., Concepcion, G.T., Feng, X., Zhang, H., Li H. (2021) Haplotype-resolved de novo assembly using phased assembly graphs with hifiasm. Nat Methods, 18:170–175. https://doi.org/10.1038/s41592-020-01056-5

Christiaens, J., F., Van Mulders, S. E., Duitama, J., et al. (2012). Functional divergence of gene duplicates through ectopic recombination. EMBO reports 13:1145–1151. https://doi.org/10.1038/embor.2012.157

Cruz-Saavedra L, Schwabl P, Vallejo GA, Carranza JC, Muñoz M, Patino LH, Paniz-Mondolfi A, Llewellyn MS, Ramírez JD. (2022). Genome plasticity driven by aneuploidy and loss of heterozygosity in Trypanosoma cruzi. Microbial Genomics 8(6): mgen000843. http://doi.org/10.1099/mgen.0.000843

Cura, C. I., Mejía-jaramillo, A. M., Duffy, T., Burgos, J. M., Rodriguero, M., Cardinal, M. V, … Schijman, A. G. (2010). Trypanosoma cruzi I genotypes in different geographical regions and transmission cycles based on a microsatellite motif of the intergenic spacer of spliced-leader genes q. International Journal for Parasitology, 40(14), 1599–1607. https://doi.org/10.1016/j.ijpara.2010.06.006

Curtis-Robles R, Hamer SA, Lane S, Levy MZ, Hamer GL. 2018. Bionomics and spatial distribution of triatomine vectors of *Trypanosoma cruzi* in Texas, USA. The American Journal of Tropical Medicine and Hygiene. 98(1): 113–121.

Danecek P, Bonfield JK, Liddle J, et al. (2021). Twelve years of SAMtools and BCFtools. GigaScience 10(2): giab008. https://doi.org/10.1093/gigascience/giab008

de Araujo CB, da Cunha JPC, Inada DT, Damasceno J, Lima ARJ, Hiraiwa P, Marques C, Gonçalves E, Nishiyama-Junior MY, McCulloch R, Elias MC. (2020). Replication origin location might contribute to genetic variability in Trypanosoma cruzi. BMC Genomics 21(1): 414. http://doi.org/10.1186/s12864-020-06803-8

de Jesus TCL, Calderano SG, Vitorino FN de L, Llanos RP, Lopes M de C, Araujo CB de, et al. (2017). Quantitative Proteomic Analysis of Replicative and Nonreplicative Forms Reveals Important Insights into Chromatin Biology of Trypanosoma cruzi. Mol Cell Proteomics 16: 23–38. https://doi.org/10.1074/mcp.M116.061200

de Lima LP, Poubel SB, Yuan ZF, Roson JN, Vitorino FNL, Holetz FB, Garcia BA, da Cunha JPC. (2020). Improvements on the quantitative analysis of Trypanosoma cruzi histone post translational modifications: study of changes in epigenetic marks through the parasite’s metacyclogenesis and life cycle. J Proteomics 225:103847. https://doi.org/10.1016/jprot.2020.103847.

de Oliveira, A. B., Chaboli, K. C., Lima, C. H., Fernanda, F., & Azeredo-Oliverira, T. V. (2018). Parasite–Vector Interaction of Chagas Disease: A Mini-Review. American Journal of Tropical Medicine and Hygiene, 98(3), 653–655. https://doi.org/10.4269/ajtmh.17-0657

De Pablos LM, Osuna A. (2012). Multigene families in Trypanosoma cruzi and their role in infectivity. Infect Immun. 80(7): 2258–2264. http://doi.org/10.1128/IAI.06225-11

Díaz-Viraqué, F., Pita, S., Greif, G., Moreira de Souza, R., Iraola, G., & Robello, C. (2019). Nanopore sequencing significantly improves genome assembly of the protozoan parasite Trypanosoma cruzi. Genome Biology and Evolution, 11(7), 1952–1957. https://doi.org/10.1093/gbe/evz129

Dujardin JC, Mannaert A, Durrant C, Cotton JA. (2014). Mosaic aneuploidy in Leishmania: the perspective of whole genome sequencing. Trends in Parasitology 30:554–555. https://doi.org/10.1016/j.pt.2014.09.004.

Echeverria, L. E., & Morillo, C. A. (2019). American Trypanosomiasis (Chagas Disease). Infectious Disease Clinics of North America, 33(1), 119–134. https://doi.org/10.1016/j.idc.2018.10.015

Elias MC, Marques-Porto R, Freymuller E, Schenkman S. Transcription rate modulation through the Trypanosoma cruzi life cycle occurs in parallel with changes in nuclear organisation. Mol Biochem Parasitol. 2001; 112: 79–90. https://doi.org/10.1016/s0166-6851(00)00349-2 PMID: 11166389

El-sayed, N. M., Myler, P. J., Bartholomeu, D. C., Nilsson, D., Aggarwal, G., Westenberger, S. J., … Vogt, C. (2005). The genome sequence of Trypanosoma cruzi, etiologic agent of Chagas disease. Science, 309, 409–415.

Falla, A., Herrera, C., Fajardo, A., Montilla, M., Adolfo, G., & Guhl, F. (2009). Haplotype identification within Trypanosoma cruzi I in Colombian isolates from several reservoirs, vectors and humans. Acta Tropica, 110(1), 15–21. https://doi.org/10.1016/j.actatropica.2008.12.003

Finn R.D., Bateman A., Clements J., Coggill P., Eberhardt R.Y., Eddy S.R., Heger A., Hetherington K., Holm L., Mistry J., et al. (2014). Pfam: the protein families database Nucleic Acids Res. 42 D222–D230

Flynn JM, Hubley R, Goubert C, Rosen J, Clark AG, Feschotte C, Smit AF. RepeatModeler2 for automated genomic discovery of transposable element families. Proc Natl Acad Sci U S A. 2020 Apr 28;117(17):9451–9457. doi: 10.1073/pnas.1921046117. Epub 2020 Apr 16. PMID: 32300014; PMCID: PMC7196820.

Franzén, O., Ochaya, S., Sherwood, E., Lewis, M. D., Llewellyn, M. S., Miles, M. A., & Andersson, B. (2011). Shotgun sequencing analysis of Trypanosoma cruzi I Sylvio X10/1 and comparison with T. cruzi VI CL Brener. PLoS Negl Trop, 5(3), 1–9. https://doi.org/10.1371/journal.pntd.0000984

Gerasimov, E. S., Zamyatnina, K. A., Matveeva, N. S., Rudenskaya, Y. A., Kraeva, N., Kolesnikov, A. A., & Yurchenko, V. (2020). Common structural patterns in the maxicircle divergent region of Trypanosomatidae. Pathogens, 9(2), 100.

Gonzalez-Garcia, L., Guevara-Barrientos, D., Lozano-Arce, D., Gil, J., Díaz-Riaño, J., Duarte, E., … & Duitama, J. (2023). New algorithms for accurate and efficient de novo genome assembly from long DNA sequencing reads. Life Science Alliance, 6(5).

Gonzalez-García, L. N., Lozano-Arce, D., Londoño, J. P., Guyot, R., & Duitama, J. (2023b). Efficient homology-based annotation of transposable elements using minimizers. Applications in Plant Sciences, e11520.

Gómez-Hernández, C., Pérez, S. D., Rezende-oliveira, K., Barbosa, C. G., Lages-silva, E., Ramírez, L. E., & Ramírez, J. D. (2019). Evaluation of the multispecies coalescent method to explore intra-Trypanosoma cruzi I relationships and genetic diversity. Parasitology, 146(8), 1063– 1074.

Greif, G., Rodriguez, M., Bontempi, I., Robello, C., & Alvarez-Valin, F. (2021) Different kinetoplast degradation patterns in American Trypanosoma vivax strains: Multiple independent origins or fast evolution?. Genomics. 113(2), 843–853. doi: 10.1016/j.ygeno.2020.12.037.

Grisard, E. C., Ribeiro, M., Paula, G., & Stoco, H. (2014). Trypanosoma cruzi Clone Dm28c Draft Genome Sequence. 2(1), 2–3. https://doi.org/10.1128/genomeA.01114-13.Copyright

Gurevich A, Saveliev V, Vyahhi N, Tesler G. QUAST: quality assessment tool for genome assemblies. Bioinformatics. 2013 Apr 15;29(8):1072–5. doi: 10.1093/bioinformatics/btt086. Epub 2013 Feb 19. PMID: 23422339; PMCID: PMC3624806.

Hernández, C., Salazar, C., Brochero, H., Teherán, A., Buitrago, L. S., Vera, M., … Ramírez, J. D. (2016). Untangling the transmission dynamics of primary and secondary vectors of Trypanosoma cruzi in Colombia: parasite infection, feeding sources and discrete typing units. Parasites & Vectors, 9(620), 1–12. https://doi.org/10.1186/s13071-016-1907-5

Herrera, C., Bargues, M. D., Fajardo, A., Montilla, M., Triana, O., Adolfo, G., & Guhl, F. (2007). Identifying four Trypanosoma cruzi I isolate haplotypes from different geographic regions in Colombia. Infection, Genetics and Evolution, 7, 535–539. https://doi.org/10.1016/j.meegid.2006.12.003

Herrera, C., Guhl, F., Falla, A., Fajardo, A., Montilla, M., Vallejo, G. A., & Bargues, M. D. (2009). Genetic Variability and Phylogenetic Relationships within Trypanosoma cruzi I Isolated in Colombia Based on Miniexon Gene Sequences. Journal of Parasitology Research, 2009(897364), 1–9. https://doi.org/10.1155/2009/897364

Jarvis, E.D., Formenti, G., Rhie, A. et al. (2022). Semi-automated assembly of high-quality diploid human reference genomes. Nature 611, 519–531. https://doi.org/10.1038/s41586-022-05325-5

Jiménez, P., Jaimes, J., Poveda, C., & Ramírez, J. D. (2019). A systematic review of the Trypanosoma cruzi genetic heterogeneity, host immune response and genetic factors as plausible drivers of chronic chagasic cardiomyopathy. Parasitology, 146(3), 269–283.

Kawashita SY, da Silva CV, Mortara RA, Burleigh BA, Briones MR. (2009). Homology, paralogy and function of DGF-1, a highly dispersed Trypanosoma cruzi specific gene family and its implications for information entropy of its encoded proteins. Molecular and Biochemical Parasitology. 165 (1):19–31. http://doi.org/10.1016/j.molbiopara.2008.12.010

Knippschild, U., Gocht, A., Wolff, S., Huber, N., Lohler, J., and Stoter, M. (2005). The casein kinase 1 family: participation in multiple cellular processes in eukaryotes. Cell Signal 17 (6), 675–689. doi: 10.1016/j.cellsig.2004.12.011

Korf I. (2004). Gene finding in novel genomes BMC Bioinform, 5 59

Krumsiek J, Arnold R, Rattei T. (2007). Gepard: A rapid and sensitive tool for creating dotplots on genome scale. Bioinformatics 23(8): 1026–1028.

Krzywinski, M., Schein, J., Birol, I., Connors, J., Gascoyne, R., Horsman, D., …& Marra, M. A. (2009). Circos: an information aesthetic for comparative genomics. Genome research, 19(9), 1639–1645.

Lander, N.; Bernal, C.; Diez, N.; Añez, N.; Docampo, R.; Ramirez, J.L. (2010). Localization and developmental regulation of a disperse gene family 1 protein in Trypanosoma cruzi. Infect. Immun. 78, 231–241.

Letunic, I. & Bork, P. (2021). Interactive Tree Of Life (iTOL) v5: an online tool for phylogenetic tree display and annotation, *Nucleic Acids Research*, Volume 49, Issue W1, W293–W296, https://doi.org/10.1093/nar/gkab301

Lewis, M. D., Llewellyn, M. S., Gaunt, M. W., Yeo, M., Carrasco, H. J., & Miles, M. A. (2009). Flow cytometric analysis and microsatellite genotyping reveal extensive DNA content variation in Trypanosoma cruzi populations and expose contrasts between natural and experimental hybrids. International journal for parasitology, 39(12), 1305–1317.jo

Li, H. (2018). Minimap2: pairwise alignment for nucleotide sequences. Bioinformatics, 34(18), 3094–3100. https://doi.org/10.1093/bioinformatics/bty191

Li L. Stoeckert C.J. Roos D.S.(2003). OrthoMCL: identification of ortholog groups for eukaryotic genomes Genome Res. 13 2178 2189

Lima ARJ, de Araujo CB, Bispo S, Patané J, Silber AM, Elias MC, et al. (2021) Nucleosome landscape reflects phenotypic differences in Trypanosoma cruzi life forms. PLoS Pathogens 17(1): e1009272. https://doi.org/10.1371/journal.ppat.1009272

Lin, R. H., Lai, D. H., Zheng, L. L., Wu, J., Lukeš, J., Hide, G., & Lun, Z. R. (2015). Analysis of the mitochondrial maxicircle of Trypanosoma lewisi, a neglected human pathogen. Parasites & vectors, 8(1), 1–11. https://doi.org/10.1186/s13071-015-1281-8.

Llewellyn, M. S., Miles, M. A., Carrasco, H. J., Lewis, M. D., Yeo, M., Vargas, J., … Gaunt, M. W. (2009). Genome-Scale Multilocus Microsatellite Typing of Trypanosoma cruzi Discrete Typing Unit I Reveals Phylogeographic Structure and Specific Genotypes Linked to Human Infection. PLoS Pathology, 5(5), 1–9. https://doi.org/10.1371/journal.ppat.1000410

Majeau, A., Murphy, L., Herrera, C., & Dumonteil, E. (2021). Assessing *Trypanosoma cruzi* Parasite Diversity through Comparative Genomics: Implications for Disease Epidemiology and Diagnostics. *Pathogens (Basel*, Switzerland), 10(2), 212. https://doi.org/10.3390/pathogens10020212

Manni, M., Berkeley, M. R., Seppey, M., & Zdobnov, E. M. (2021). BUSCO: assessing genomic data quality and beyond. Current Protocols, 1(12), e323.

Manoel-Caetano, F. da S., & Silvia, A. E. (2007). Implications of genetic variability of Trypanosoma cruzi for the pathogenesis of Chagas disease. Cadernos de Saúde Pública, 23(10), 2263–2274.

Marcili, A., Lima, L., Cavazzana, M., Junqueira, A. C. V, Veludo, H. H., Da Silva, F. M., … Teixeira, M. M. G. (2009). A new genotype of Trypanosoma cruzi associated with bats evidenced by phylogenetic analyses using SSU rDNA, cytochrome b and Histone H2B genes and genotyping based on ITS1 rDNA A new genotype of Trypanosoma cruzi associated with bats evidenced by phyloge. Parasitology, 136(6), 641–655. https://doi.org/10.1017/S0031182009005861

Mejía-Jaramillo, A. M., Peña, V. H., & Triana-Chávez, O. (2009). Trypanosoma cruzi: biological characterization of lineages I and II supports the predominance of lineage I in Colombia. Experimental parasitology, 121(1), 83–91.

Ministerio de Salud y Protección Social. (2010). Guía Protocolo para la vigilancia en salud pública de Chagas. Bogotá. Instituto Nacional de Salud, 7. https://www.minsalud.gov.co/Documents/Salud%20P%C3%BAblica/Ola%20invernal/Protocolo%20Chagas.pdf

Minning, T. A., Weatherly, D. B., Flibotte, S., & Tarleton, R. L. (2011). Widespread, focal copy number variations (CNV) and whole chromosome aneuploidies in Trypanosoma cruzi strains revealed by array comparative genomic hybridization. BMC Genomics, 12(139), 1–11.

Moraes Barros RR, Marini MM, Antonio CR, Cortez DR, Miyake AM, Lima FM, et al. (2012). Anatomy and evolution of telomeric and subtelomeric regions in the human protozoan parasite Trypanosoma cruzi. BMC Genomics 13: 229.

Myler, P. J., Glick, D., Feagin, J. E., Morales, T. H., & Stuart, K. D. (1993). Structural organization of the maxicircle variable region of Trypanosoma brucei: identification of potential replication origins and topoisomerase II binding sites. Nucleic acids research, 21(3), 687–694.

Organización Panamericana de Salud (n.d). Enfermedad de Chagas. https://www.paho.org/es/temas/enfermedad-chagas

Otto T.D. Dillon G.P. Degrave W.S. Berriman M. RATT: Rapid Annotation Transfer Tool Nucleic Acids Res.2011 39 e57

Rachidi N, Taly JF, Durieu E, et al., “Pharmacological assessment defines Leishmania donovani casein kinase 1 as a drug target and reveals important functions in parasite viability and intracellular infection,” Antimicrobial Agents and Chemotherapy, vol. 58, no. 3, pp. 1501–1515, 2014.

Ramírez, J. D., Guhl, F., Rendón, L. M., Rosas, F., Marin-neto, J. A., & Morillo, C. A. (2010). Chagas Cardiomyopathy Manifestations and Trypanosoma cruzi Genotypes Circulating in Chronic Chagasic Patients. PLoS Neglected Tropical Diseases, 4(11), 1–9. https://doi.org/10.1371/journal.pntd.0000899

Ramírez, J. D., Tapia-calle, G., & Guhl, F. (2013). Genetic structure of Trypanosoma cruzi in Colombia revealed by a High-throughput Nuclear Multilocus Sequence Typing (nMLST) approach. BMC Genetics, 14(96).

Rassi, A. J., Rassi, A., & De Rezende, J. M. (2012). American Trypanosomiasis (Chagas Disease). Infectious Disease Clinics of North America, 26, 275–291. https://doi.org/10.1016/j.idc.2012.03.002

Reis-cunha, J. L., Rodrigues-Luiz, G. F., Valdivia, H. O., Baptista, R. P., Mendes, T. A. O., Morais, G. L. De, … Bartholomeu, D. C. (2015). Chromosomal copy number variation reveals differential levels of genomic plasticity in distinct Trypanosoma cruzi strains. BMC Genomics, 16(499), 1–15. https://doi.org/10.1186/s12864-015-1680-4

Reis-Cunha, J.L., Valdivia, H.O., Bartholomeu, D.C. (2018). Gene and Chromosomal Copy Number Variations as an Adaptive Mechanism Towards a Parasitic Lifestyle in Trypanosomatids. Current Genomics 19, 87–97.

Rice, P., Longden, I., & Bleasby, A. (2000). EMBOSS: the European Molecular Biology Open Software Suite. Trends in genetics : TIG, 16(6), 276–277. https://doi.org/10.1016/s0168-9525(00)02024-2

Rosón JN, Vitarelli MO, Costa-Silva HM, Pereira KS, Pires DDS, Lopes LS, Cordeiro B, Kraus AJ, Cruz KNT, Calderano SG, Fragoso SP, Siegel TN, Elias MC, da Cunha JPC. H2B.V demarcates divergent strand-switch regions, some tDNA loci, and genome compartments in Trypanosoma cruzi and affects parasite differentiation and host cell invasion. PLoS Pathog. 2022 Feb 18;18(2):e1009694. doi: 10.1371/journal.ppat.1009694

Ruvalcaba-Trejo, L. I. & Sturm, N. R. (2011). The Trypanosoma cruzi Sylvio X10 strain maxicircle sequence: the third musketeer. BMC Genomics. 12:58. doi: 10.1186/1471-2164-12-58

Schmunis, G. A., & Yadon, Z. E. (2010). Chagas disease: A Latin American health problem becoming a world health problem. Acta Tropica, 115(1–2), 14–21. https://doi.org/10.1016/j.actatropica.2009.11.003

Simpson, L., Neckelmann, N., de La Cruz, V. F., Simpson, A. M., Feagin, J. E., Jasmer, D. P., & Stuart, J. E. (1987). Comparison of the maxicircle (mitochondrial) genomes of Leishmania tarentolae and Trypanosoma brucei at the level of nucleotide sequence. Journal of Biological Chemistry, 262(13), 6182–6196.

Souto, R. P., Fernandes, O., Macedo, A. M., Campbell, D. A., & Zingales, B. (1996). DNA markers define two major phylogenetic lineages of Trypanosoma cruzi. Molecular and biochemical parasitology, 83(2), 141–152.

Souza, R. T., Lima, F. M., Barros, R. M., Cortez, D. R., Santos, M. F., Cordero, E. M., … Da Silveira, J. F. (2011). Genome Size, Karyotype Polymorphism and Chromosomal Evolution in Trypanosoma cruzi. PLoS ONE, 6(8), 1–14. https://doi.org/10.1371/journal.pone.0023042

Spadafora C, Repetto Y, Torres C, Pino L, Robello C, Morello A, Gamarro F, Castanys S. Two casein kinase 1 isoforms are differentially expressed in Trypanosoma cruzi. Mol Biochem Parasitol. 2002 Sep-Oct;124(1-2):23–36. doi: 10.1016/s0166-6851(02)00156-1. PMID: 12387847.

Stanke M. Schöffmann O. Morgenstern B. Waack S. (2006). Gene prediction in eukaryotes with a generalized hidden Markov model that uses hints from external sources BMC Bioinform. 7 62

Steinbiss, S., Silva-franco, F., Brunk, B., Foth, B., Hertz-fowler, C., Berriman, M., & Otto, T. D. (2016). Companion : a web server for annotation and analysis of parasite genomes. Nucleic Acids Research, 44(1), 29–34. https://doi.org/10.1093/nar/gkw292

Sturm, N. R., Vargas, N. S., Westenberger, S. J., Zingales, B., & Campbell, D. A. (2003). Evidence for multiple hybrid groups in Trypanosoma cruzi. International Journal for Parasitology, 33(3), 269– 279.

Talavera-López, C., Messenger, L. A., Lewis, M. D., Yeo, M., Reis-Cunha, J. L., Matos, G. M., … & Andersson, B. (2021). Repeat-driven generation of antigenic diversity in a major human pathogen, Trypanosoma cruzi. Frontiers in cellular and infection microbiology, 11, 614665.

Tello D, Gonzalez-Garcia LN, Gomez J, Zuluaga-Monares JC, Garcia R, Angel R, Mahecha D, Duarte E, Leon MDR, Reyes F, Escobar-Velásquez C, Linares-Vásquez M, Cardozo N, Duitama J. (2023). NGSEP 4: Efficient and accurate identification of orthogroups and whole-genome alignment. Mol Ecol Resour. Apr;23(3):712–724. doi: 10.1111/1755-0998.13737. Epub 2022 Nov 27. PMID: 36377253.

Tibayrenc, M. (1998). Genetic epidemiology of parasitic protozoa and other infectious agents: the need for an integrated approach. Int J Parasitol, 28(1), 85–104.

Triana O, Ortiz S, Dujardin JC, Solari A. 2006. Trypanosoma cruzi: Variability of stock from Colombia determined by molecular karyotype and minicircle Southern blot analysis. Experimental Parasitology, 113; 62–66.

Urbaniak M.D., “Casein kinase 1 isoform 2 is essential for bloodstream form Trypanosoma brucei,” Molecular and Biochemical Parasitology, vol. 166, no. 2, pp. 183–185, 2009.

Urrea, D. A., Duitama, J., Imamura, H., Alzate, J. F., Gil, J., Muñoz, N., … Triana-Chavez, O. (2018). Genomic Analysis of Colombian Leishmania panamensis strains with different level of virulence. Scientific Reports, 1–16. https://doi.org/10.1038/s41598-018-35778-6

Urrea D.A., Triana-Chavez O., Alzate J.F. (2019). Mitochondrial genomics of human pathogenic parasite Leishmania (Viannia) panamensis. PeerJ 7:e7235. http://doi.org/10.7717/peerj.7235

Vallejo, G. A., Guhl, F., Carranza, J. C., Lozano, L. E., Sánchez, J. L., Jaramillo, J. C., … & Steindel, M. (2002). kDNA markers define two major Trypanosoma rangeli lineages in Latin-America. Acta Tropica, 81(1), 77–82.

Villa LM, Guhl F, Zabala D, Ramírez JD, Urrea DA, Diana Carolina Hernández, Zulma Cucunubá, Marleny Montilla, Julio César Carranza, Karina Rueda, Jorge Eduardo Trujillo, Gustavo Adolfo Vallejo (2013). The identification of two Trypanosoma cruzi I genotypes from domestic and sylvatic transmission cycles in Colombia based on a single polymerase chain reaction amplification of the spliced-leader intergenic region. Mem Inst Oswaldo Cruz, Rio de Janeiro, Vol. 108(7): 932–935

Wang W, Peng D, Baptista RP, Li Y, Kissinger JC, Tarleton RL (2021) Strain-specific genome evolution in *Trypanosoma cruzi*, the agent of Chagas disease. PLoS Pathog 17(1): e1009254. https://doi.org/10.1371/journal.ppat.1009254

Westenberger, S. J., Barnabé, C., Campbell, D. A., & Sturm, N. R. (2005). Two Hybridization Events Define the Population Structure of Trypanosoma cruzi. Genetics, 171(2), 527–543. https://doi.org/10.1534/genetics.104.038745

World Health Organization. (2020). Chagas disease (also known as American trypanosomiasis). Retrieved from https://www.who.int/en/news-room/fact-sheets/detail/chagas-disease-(american-trypanosomiasis)

Yang, Z. (2007). PAML 4: a program package for phylogenetic analysis by maximum likelihood. Molecular Biology and Evolution 24: 1586–1591.

Zingales, B, Andrade, S. G., Briones, M. R. S., Campbell, D. A., Chiari, E., Fernandes, O., … Schijman, A. G. (2009). A new consensus for Trypanosoma cruzi intraspecific nomenclature: second revision meeting recommends TcI to TcVI. Memórias Do Instituto Oswaldo Cruz, 104(7), 1051–1054.

Zingales, Bianca, Miles, M. A., Campbell, D. A., Tibayrenc, M., Macedo, A. M., Teixeira, M. M. G., … Sturm, N. R. (2012). Infection, Genetics and Evolution The revised Trypanosoma cruzi subspecific nomenclature: Rationale, epidemiological relevance and research applications. Infection, Genetics and Evolution, 12(2), 240–253. https://doi.org/10.1016/j.meegid.2011.12.009

Zingales B. (2018). *Trypanosoma cruzi* genetic diversity: Something new for something known about Chagas disease manifestations, serodiagnosis and drug sensitivity. Acta Trop 184:38–52

